# Dysfunction of the episodic memory network in the Alzheimer’s disease cascade

**DOI:** 10.1101/2024.10.25.620237

**Authors:** René Lattmann-Grefe, Niklas Vockert, Judith Machts, Yanin Suksangkharn, Renat Yakupov, Hartmut Schütze, Wenzel Glanz, Enise Incesoy, Michaela Butryn, Falk Lüsebrink, Matthias Schmid, Melina Stark, Luca Kleineidam, Annika Spottke, Marie Coenjaerts, Frederic Brosseron, Klaus Fliessbach, Anja Schneider, Peter Dechent, Stefan Hetzer, Klaus Scheffler, Alfredo Ramirez, Christoph Laske, Sebastian Sodenkamp, Slawek Altenstein, Luisa-Sophie Schneider, Daria Gref, Eike Jakob Spruth, Andrea Lohse, Björn H. Schott, Jens Wiltfang, Ingo Kilimann, Doreen Goerss, Ayda Rostamzadeh, Josef Priller, Oliver Peters, Julian Hellmann-Regen, Stefan Teipel, Frank Jessen, Anne Maass, Gabriel Ziegler, Emrah Düzel

## Abstract

Alzheimer’s disease (AD) is a major cause of dementia and cognitive decline. Here we assessed how episodic memory circuit dysfunction, a hallmark of AD, is related to the longitudinal cascade of AD biomarkers, neurodegeneration and cognition using data from the DZNE Longitudinal Cognitive Impairment and Dementia study. This data set is unique by including over 1000 longitudinal functional magnetic resonance imaging (fMRI) measurements during episodic memory encoding. We leveraged a disease progression model (DPM) to obtain AD progression scores. Voxel-wise analyses revealed widespread loss of deactivation (hyperactivation) and activation (hypoactivation) with increasing disease stage. Hyperactivation trajectories were nonlinear and visually preceded trajectories of cognition. Overall, hyperactivation was independently associated with co-occurrence of amyloid- and tau-positivity and neurodegeneration, suggesting synaptic dysfunction and neurodegeneration as two independent drives of cognitive decline. Our results therefore provide evidence for a critical time window in which pharmacological treatments targeting the synapse may improve cognition.

---

Alzheimer’s disease (AD) is one of the leading causes of dementia and cognitive decline in Western societies^1^. Cognitive decline, and particularly memory loss, is the predominant clinical symptom of AD and the primary endpoint of disease-modifying, symptomatic and risk-reducing treatments^2^. A major challenge is to uncover how cognitive decline relates to the progression of AD. According to the amyloid-cascade hypothesis^3^, cognitive decline is the end-stage of a progression from Aβ aggregation to accelerated spreading of hyperphosphorylated tau and consequently to neurodegeneration. However, animal models indicate that cognitive dysfunction can precede neurodegeneration as a result of pre- and postsynaptic dysfunction due amyloid and tau pathology^4–7^. Here, we addressed this discrepancy leveraging a longitudinal disease progression model (DPM) of human AD for the first time in conjunction with measures of cognitive brain activity as a proxy for synaptic dysfunction^8^. This allowed us to model the relationship between cognitive decline and brain activity abnormalities in relation to pathological hallmarks of AD. We utilized cross-sectional and longitudinal measures of CSF- Aβ42/40 ratios, CSF-phosphotau181 (pTau181), and neurodegeneration (i.e., volume loss) along the amyloid cascade.

According to an influential model^9^, the change towards abnormality in AD biomarkers follows a sigmoidal, monotonically increasing disease progression trajectory, where cognitive decline follows neurodegeneration. The unfolding of this hypothetical cascade over up to two decades^10^ has been captured within the ATN framework^11^. Investigation of such models requires the conjoint availability of longitudinal CSF biomarkers, volumetry and cognitive markers across the whole AD spectrum ranging from cognitively unimpaired to mild cognitive impairment and mild dementia. This is highlighted by recent advances from studies using continuous DPMs. Results showed that time frames spanning the whole disease cascade could be estimated from shorter-scale longitudinal data^12–15^.

Episodic memory, the ability to recall recent personal experiences^16,17^, is critically dependent on the medial temporal lobe (MTL) memory system and structures of the so-called Default Model Network (DMN)^18^. It is one of the first cognitive faculties to be impaired along the AD cascade^2^. Consequently, a major effort of therapeutic and interventional studies is to ameliorate episodic memory decline in AD through interventions, including lifestyle^19^, pharmacology^20^ or transcranial brain stimulation^21^. Recent data from anti-amyloid antibody trials indicate that, in early AD, amyloid antibodies may improve cognition^22,23^. However, this observation is not compatible with a cascade model in which cognitive dysfunction follows neurodegeneration but rather suggests a major contribution of synaptic dysfunction to cognitive impairment^7^. Thus, uncovering the dynamics of episodic memory circuitry dysfunction across AD progression may aid at improving individualized treatment for patients with AD.

Brain activity related to episodic memory can be reliably measured using task-based functional MRI (fMRI). Task fMRI has been extensively used to investigate episodic memory both in healthy individuals and individuals affected by AD^8,24–32^. Here, we used task-based fMRI as a proxy for synaptic dysfunction in the human episodic memory network. We defined synaptic dysfunction as a brain activity abnormality that cannot be explained by neurodegeneration^8^. This definition follows the rationale from animal research, where synaptic dysfunction is conceptualized independently from synapse loss^5^. In line with this rationale, neurons in the vicinity of soluble and insoluble Aβ deposits in AD mouse models show hyperactivation compared to neurons distant from deposits in the hippocampus^33,34^. In the face of combined Aβ- and tau pathology, on the other hand, neurons show hypoactivation^5^. Hyper- and hypoactivation-like patterns have also been observed in human task fMRI studies of AD. These studies have reported lower deactivations in regions normally deactivated (also denoted as “hyperactivation” and “reduced deactivation”, respectively) during memory tasks like novelty detection or successful encoding, most notably structures of the DMN like the posterior cingulate cortex and precuneus^26,35–40^. For areas activated rather than deactivated during memory tasks (e.g., the inferior temporal lobe and hippocampus), a number of studies have reported hypoactivation^8,26,27^. Our study builds on these reported brain activation abnormalities to model their relationship to AD progression in a DPM.

Here, we generated a computational disease progression marker using a multivariate DPM approach and investigated its relationship with episodic memory-related task-fMRI activation using the largest to date available longitudinal task-fMRI data set in AD. We studied longitudinal changes in task-fMRI activation patterns over multiple follow-ups. Our analyses were performed in the DELCODE study that covers the full pre-clinical and clinical AD spectrum, including cognitively normal older adults (CN), individuals with subjective cognitive decline (SCD) and patients with MCI and mild dementia of the Alzheimer’s type (DAT)^41^. We hypothesized that there would be region-specific associations between encoding-related brain activity and latent disease progression stage. Additionally, we hypothesized that AD-related brain activity patterns and brain activity changes would in part indicate synaptic dysfunction by being independent of neurodegeneration.

## 2. Results

### 2.1 Disease stage associates with several AD-related factors

Baseline demographics are displayed in Supplementary Table 1. In order to obtain a continuous AD-related disease stage score, we used 739 longitudinal ATN biomarker and cognitive measurements from 208 participants to train the probabilistic DP model (see methods for details; Supplementary Table 2 for baseline characteristics of this subsample). The results of this model were two-fold. First, empirical biomarker disease progression curves (Fig. 1A, Supplementary Figure 1) were obtained. Second, each participant was assigned a continuous disease stage value on an arbitrary scale in years. Note that obtained disease stages are probabilistic estimates (given the ATN and cognitive test score of the patient), characterizing relative rather than absolute stages reflecting individual differences in the analyzed sample. Bivariate correlations between DPM markers and obtained disease stages are provided in Supplementary Figure 2. Inspecting timepoints of fastest changes, we noted that CSF biomarkers became abnormal first, followed by brain volume estimates and, ultimately, cognitive performance (Fig. 1B). Investigating the associations of our disease stage scores with demographics in a multiple regression analysis, we found demographics to contribute significantly to the disease stage variability (F(3,489) = 42.56, *p* < .001, adjusted R^2^ = 0.202). Specifically, higher age was indicative of a more advanced disease stage (t = 9.148, *p <* .001), whereas higher education was related to earlier disease stages (t = −4.979, *p <* .001). Sex showed no significant association with estimated disease stage (t = 0.637, *p* = .524). All further analyses with disease stage consider the age-, sex-, and education-corrected disease stage values.

**Figure 1.**
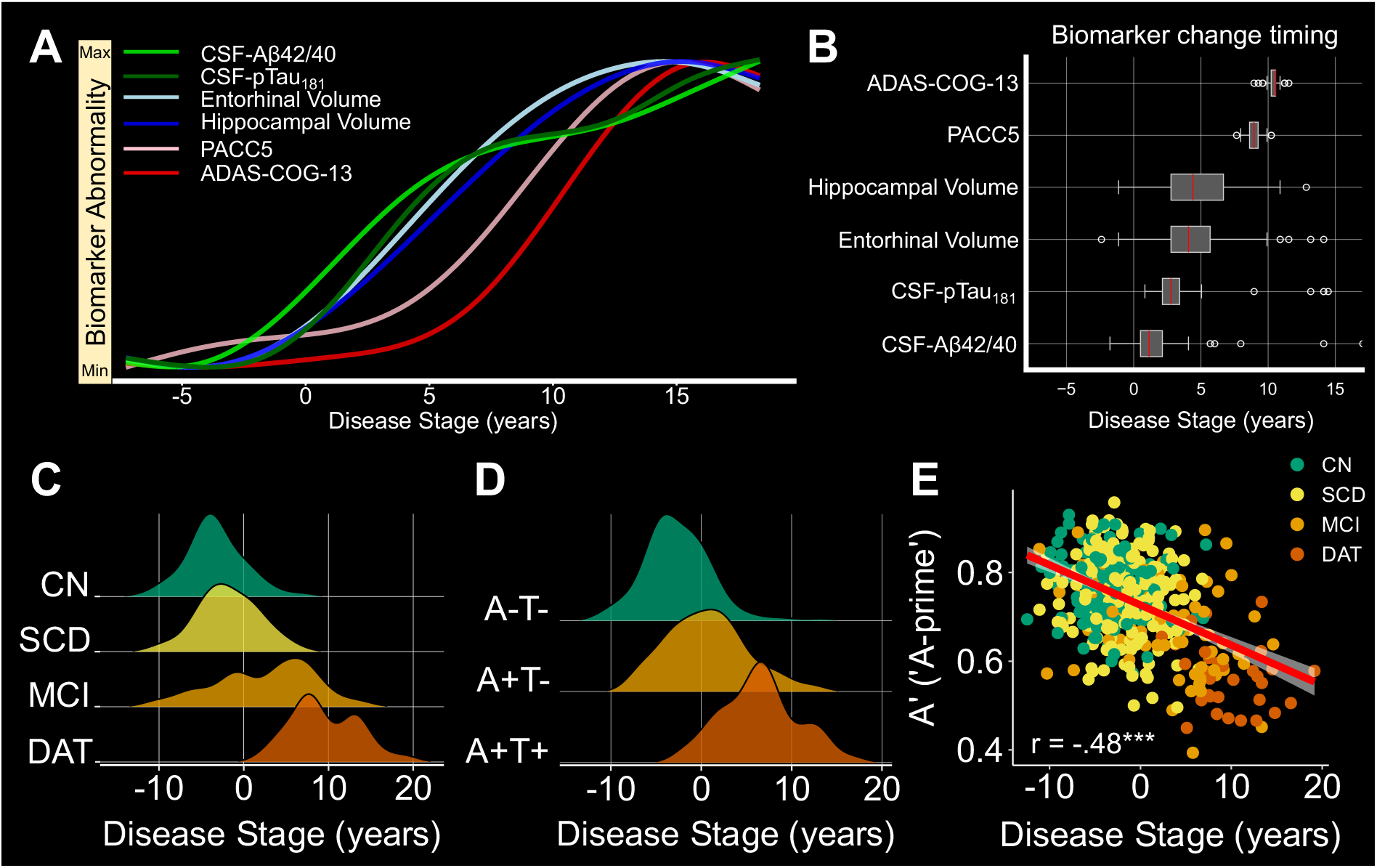
Disease progression curves and association with AD-related variables. **A** GP-based DPM used in this study comprising longitudinal CSF (Aß42/40, pTau181), volumetric MRI (hippocampal, entorhinal volume), and PACC5 as well as ADAS-COG-13 cognitive score within the DELCODE cohort using 787 available data points from 210 participants (82 CN, 57 A+ SCD, 44 A+ MCI, 27 A+ DAT). **B** Timepoints of fastest changes derived from temporal derivatives of the empirical biomarker progression curves from the model posterior. The timepoint of fastest change was sampling 200 times from the model posterior and the red line indicates its median. **C** Ridgeline plots of disease stages in diagnostic groups. **D** Ridgeline plots of disease stages in AT classification subgroups of AD pathology. **E** Associations between the estimated disease stage and memory performance during the task-fMRI session residualized for age, sex, and education. \*\*\**p* < .001.

Next, associations with diagnostic groups and AT biomarker categories were investigated. Diagnostic group membership was significantly related to disease stage (F(3,479) = 133.11, *p <* .001), reflecting the *a priori* assignment of participants to diagnostic groups at baseline. *Post-hoc* analysis revealed the following ordering of disease stage for baseline diagnosis: CN < SCD < MCI < DAT (Fig. 1C; Supplementary Table 3). Additionally, when analyzing disease stage differences over the AT criteria, biomarker groups were significantly associated with disease stage (F(2,209) = 92.11, *p* < .001). Ordering of disease stages by AT groups was as follows: A- T- < A+T- < A+T+ (Fig. 1D, Supplementary Table 4).

Lastly, we were interested in the association between disease stages and a fMRI task performance marker not used in model fitting. As expected, participants further in disease progression showed worse A’ scores of fMRI task performance (r = -.48, *p* < .001; Fig. 1E) accounting for age, sex, and education.

### 2.2 Encoding-related brain activity changes with progression towards AD

After quantitatively establishing a marker of a continuous disease score characterizing progression towards AD in the previous section, we investigated encoding-related activity differences as a function of disease progression. Such differences during the course of AD are potentially reflective of disease-related functional reorganization and adaptation. In the whole sample (n = 493), successful memory encoding was related to activation of a large network including lingual gyri, occipital, and prefrontal regions, while widespread deactivations were observed in the precuneus, posterior cingulate cortex, inferior parietal lobule, and fronto-temporal regions (Fig. 2A), replicating previous observations in the same dataset^39,40,42^ and in independent cohorts^35,37,38,43^. We observed that more advanced disease scores were related to hyperactivation in the precuneus, inferior parietal lobule, and posterior cingulate cortices bilaterally, as well as anterior cingulate cortex and superior frontal gyrus (Fig. 2B, orange colors, Family-Wise error (FWE) corrected), reflecting similar previous observations based on the novelty contrast of the same paradigm^26^. Additionally, we found right-lateralized reductions in encoding-related activation with advancing disease progression within the middle occipital gyrus and inferior temporal gyrus (Fig. 2B, blue colors, FWE corrected). Although we tested for non-linearities, we did not find indications for inverted U-shaped associations between disease stage and successful memory encoding. In summary, we found hyperactivations (or reduced deactivations) as well as a reduction in activation during successful memory encoding with further disease progression.

**Figure 2.**
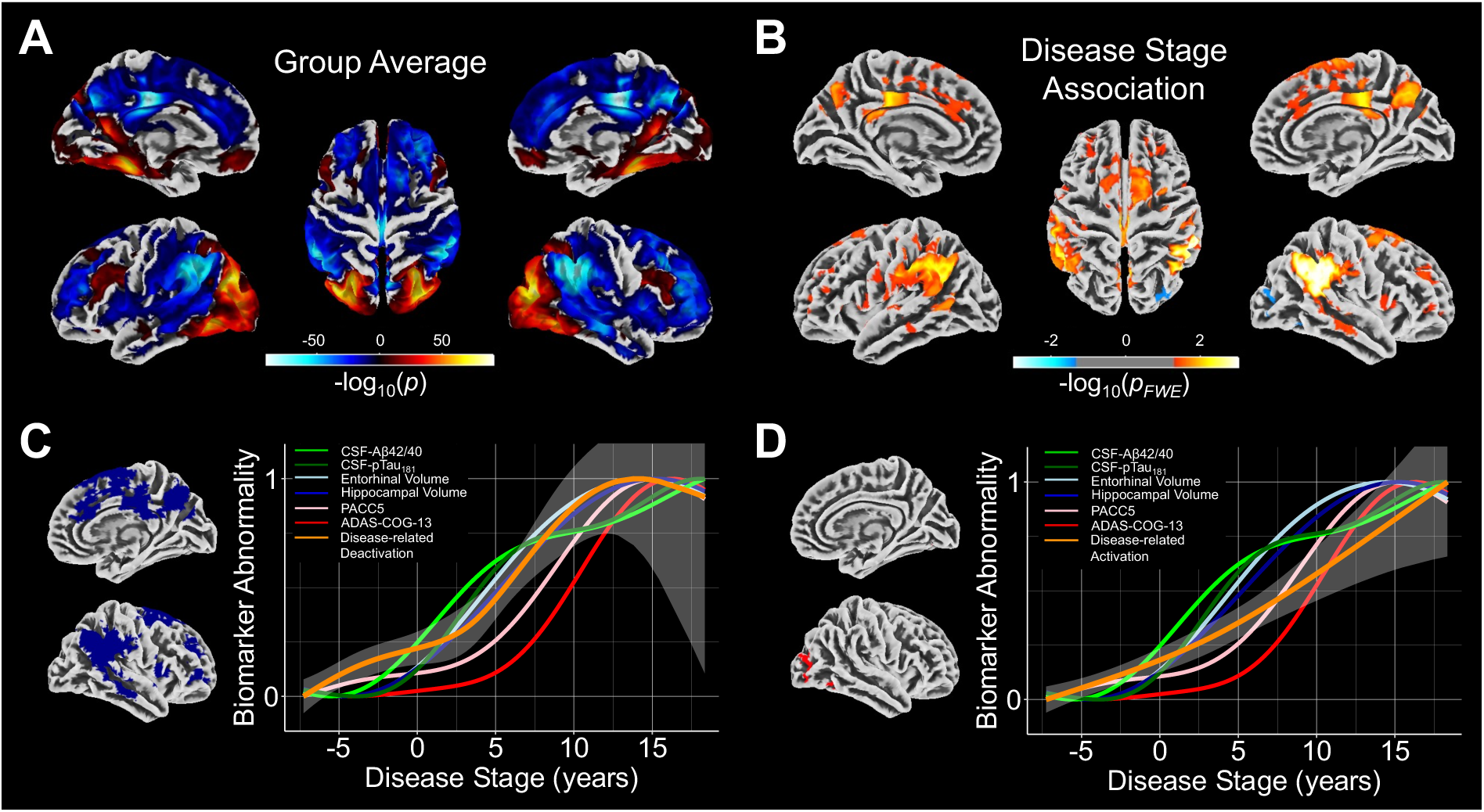
Association between encoding-related activity and disease stages. **A** Surface representation of the average activation (red colors) and deactivation (blue colors) for successful memory encoding extracted from the second-level SwE model (n = 493). **B** Linear associations of disease stage with successful memory encoding related activation (red colors) and deactivation (blue colors) from the second-level SwE model. Results were obtained using the Wild bootstrap method with 1000 repetitions. **C** Relative position of biomarker abnormality differences for encoding-related deactivation in relation to the biomarkers from our data-driven DPM approach. FMRI abnormality starts in the earliest disease stages. Towards disease stage 0, a small plateau can be observed before increasing in abnormality together with volumetric biomarkers. **D** Relative position of biomarker abnormality change for the successful memory encoding related activation. For activation, we only observed a linear increase in abnormality over the disease progression.

### 2.3 Towards task-activity based AD biomarker progression curves

Since the progression of activity changes in patients transitioning towards AD exhibited unexpected dynamics (e.g. in early stages), we next assessed activity progression curves using a more flexible curve fitting approach. Obtained progression trajectories from mean activity in clusters from the above analyses are presented in Fig. 2C and D. Deactivation followed a non-linear progression trajectory, which was characterized by an initial increase of abnormality in the earliest disease stages, followed by the most pronounced changes in later disease stages (F_non-linear_(2,489) = 3.793, *p*_uncorr_ = .023, *p*_Holm_ = .046; Fig. 2C). These later changes were accompanied by changes in hippocampal and entorhinal cortex volume. Cognition started to show abnormalities after memory-related fMRI deactivations (time points of fastest change (in years) - Aβ42/40: 1.17; pTau_181_: 2.79; Entorhinal Volume: 4.09; Hippocampal Volume: 5.38; Episodic Memory Deactivation: 6.36; PACC5: 8.95; ADAS-COG-13: 10.57; Fig. 1B). Disease-related changes in fMRI activations, on the other hand, resembled a monotonic and largely linear increase (Fig. 2D).

### 2.4 Associations between subsequent memory and individual biomarkers from the DPM

The disease stage is a marginal score combining the influence of several components into one single index. Thus, we were further interested in individual contributions of biomarkers towards activity variability beyond demographic information (age, sex, and years of education). First, AT group membership was significantly associated with memory-related deactivation (F(2,157) = 13.35, *p* < .001, η^2^_partial_ = .15 [95% CI: .07; 1]). *Post-hoc* tests revealed that combined amyloid and tau positivity (A+T+) was related to reduced deactivations compared to the other groups (A-T-, *t* = 4.981, *p* < .001, and A+T-, *t* = 4.397, *p* < .001). A correction for hippocampal and entorhinal volume (F(2,155) = 4.82, *p* = .009, η^2^_partial_ = .06 [.01; 1]) did not change the association (A-T-, *t* = 2.81, *p_Holm_* = .007, and A+T-, *t* = 2.46, *p_Holm_* = .022; Fig. 3A, blue colors). An additional correction for both volumes and both cognitive measures from our DPM (F(2,153) = 3.12, *p* = .047, η^2^_partial_ = .04 [.0003; 1]) did not change the association. For memory-related fMRI activation (task-positive regions), we also found a significant association with AT group membership (F(2,157) = 6.212, *p* = .003, η^2^ = .07 [.02; 1]). *Post-hoc* tests revealed that this was driven by the lower activation in the A+T+ compared to the A-T- group (*t* = −3.465, *p* = .002). While additional control for hippocampal and entorhinal volume did not change the association, (F(2,155) = 4.31, *p* = .015, η^2^_partial_ = .05 [.01; 1]; *Post-hoc* analysis (A- T-), t = −2.14, *p_Holm_* = .012, Fig. 3A, red colors), AT group membership was not related to variability in successful memory-related activation when controlling for volume and cognition measures from our DPM combined (F(2,153) = 1.56, *p* = .213, η^2^_partial_ = .02 [0; 1]).

**Figure 3.**
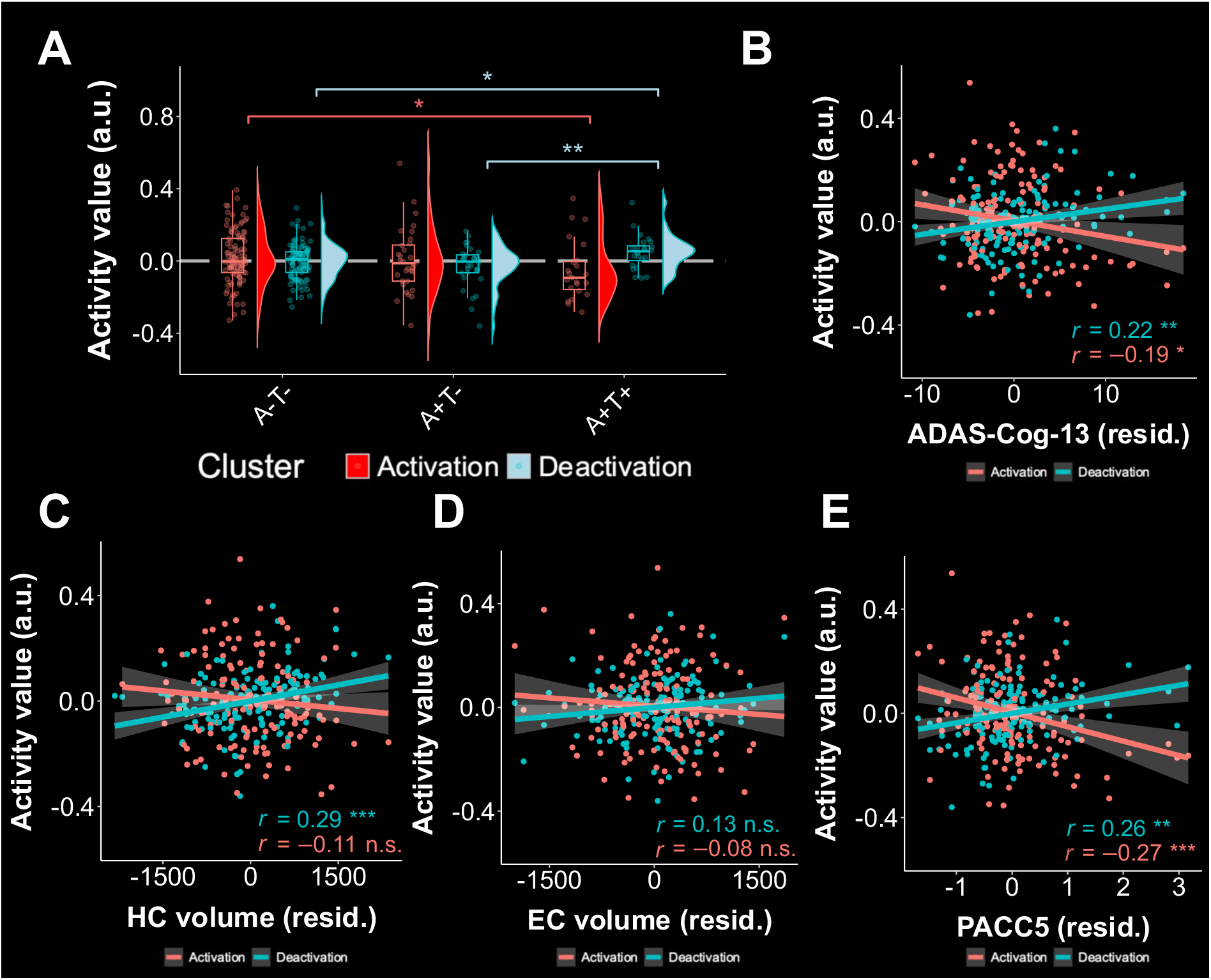
Distinct contributions of DPM biomarkers to disease-related activation variability in successful memory encoding. A-E We used a subsample of n = 160 individuals (n_A-T-_ = 104, n_A+T-_ = 32, n_A+T+_ = 24) with available fMRI and complete biomarker data (i.e. CSF, volume, and cognition) to determine unique contributions of DPM biomarker groups to disease-related activation variability. **A** Association between AD biomarker staging groups and encoding-related activity, corrected for volume and cognition differences. **B-E** Semi-partial correlations between activity and biomarkers of atrophy and cognition. Each group of biomarkers is corrected for the influence of the other two, respectively. All variables are additionally corrected for age, sex, and education.

Next, we calculated semi-partial correlation coefficients for the association of DPM biomarkers with activation and deactivation values, correcting for the influences of other biomarkers in the DPM, respectively (Fig. 3B-E). For this, we multiplied volumes and the PACC5 score by −1, such that higher values in all DPM variables indicate higher amounts of pathology. We observed that, when controlling for CSF biomarkers, hippocampal volume was positively associated with mean cluster deactivation (r = .29, *p* < .001, Fig. 3C) but not with activation (r = -.11, *p* = .183, Fig. 3C). Entorhinal cortex atrophy was not correlated with fMRI activity (all p > .05, Fig. 3D). Regarding cognitive performance, we found that encoding-related deactivation was significantly positively correlated with PACC5 (r = .26, *p* < .001) and with ADAS-COG-13 (r = .22, *p* = .009) when the influence of hippocampal and entorhinal volumes were controlled for. Encoding-related activation was significantly negatively correlated with PACC5 scores (r = -.27, *p* < .001), and with ADAS-COG-13 (r = -.19, *p* = .016) when controlling for both volumes. In summary, the emergence of tau and hippocampal volume provide unique explanatory variance to disease-related fMRI activation differences. Additionally, beyond the influences of volume, brain activity explained cognitive performance differences.

### 2.5 Longitudinal activation change is moderated by disease stage

If longitudinal data were reflective of our empirical disease curves for fMRI from above, this would provide further leverage for establishing fMRI as a biomarker in AD. To this end, we hypothesized that the disease stage variable would moderate longitudinal changes in successful encoding-related activity across follow-ups, particularly in deactivation. Random-intercept-random-slope models did not converge, which is why we report on the random intercept models only. The corresponding tables can be found in the supplement (Supplementary Tables 5 and 6, respectively). As expected, we found change over follow-ups in encoding-related deactivation to be significantly moderated by disease stage (*t*(322.380) = 2.436, *p*_uncorrected_ = .01540, *p*_Bonferroni-Holm_ = .0308, *η^2^*_partial_ = .019 [.00186;1]; Fig. 4A). On the other hand, this was not the case for encoding-related activations (*t*(317.37) = 1.659, *p*_uncorrected_ = .098, *p*_Bonferroni-Holm_ = .098, *η^2^*_partial_ = .008 [0;1]; Fig. 4B). Finally, we investigated associations between longitudinal changes of both encoding-related fMRI activation and disease progression. To this end, we correlated individual-level activation and deactivation slopes over follow-ups with intra-individual slopes of the disease stage variable. Neither changes in activation nor in deactivation were significantly correlated with the changes in disease stage in our sample (all *p* > 0.05).

**Figure 4.**
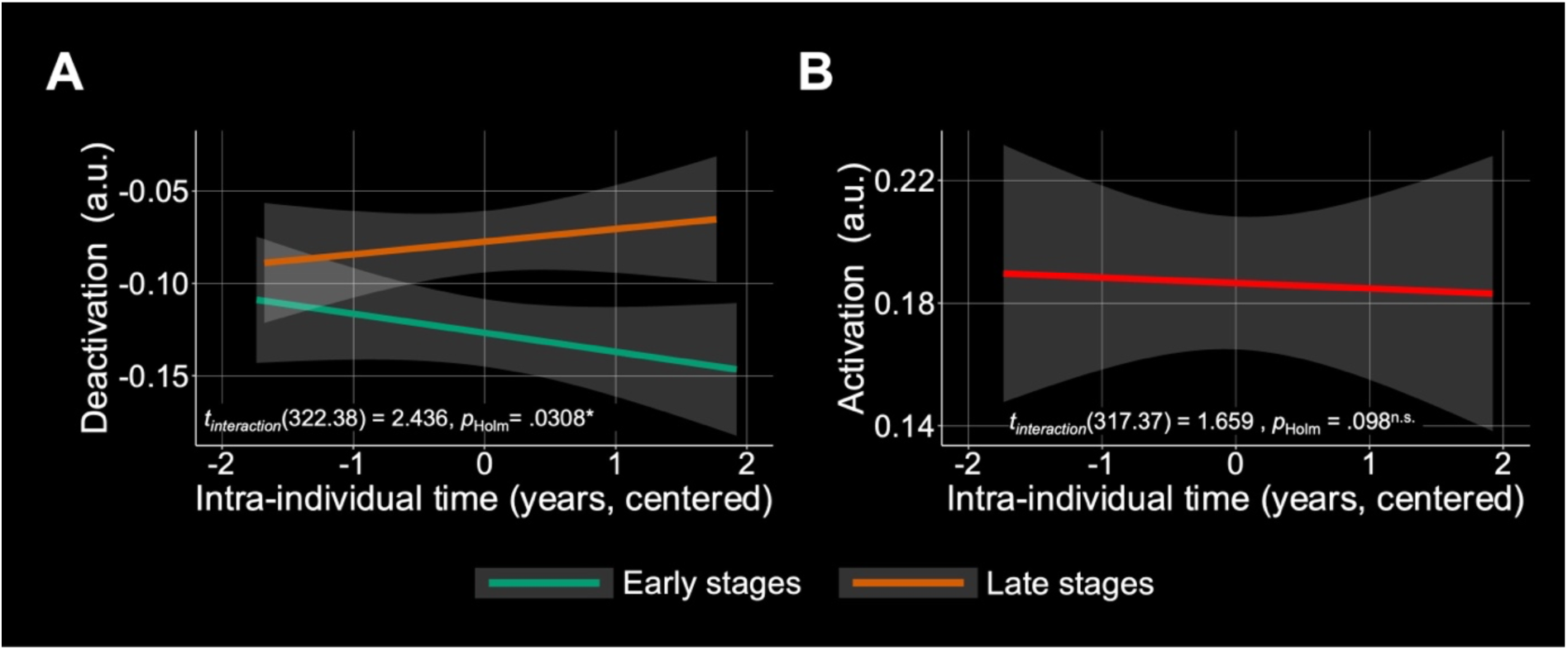
Association between disease stage and longitudinal change of successful memory encoding–related activation and deactivation. For this analysis, longitudinal fMRI data from n = 166 individuals with available AD biomarker information was used (n_A-T-_ = 108, n_A+T-_ = 33, n_A+T+_ = 27). The continuous disease stage scores were used to group participants into early and late stages based on the median disease stage (−3.37) in the sample. **A** Interaction effect Time x Disease Stage on disease-related encoding deactivation. **B** The same interaction effect as tested in A, but for disease-related encoding activation. Here, change over follow-ups is not significantly moderated by disease stage. Thus, we report the overall slope for the whole sample over follow-ups. P-values were corrected using the Bonferroni-Holm procedure. * *p*_Holm_ < .05; ^n.s.^ *p*_Holm_ > .05.

## 3. Discussion

In the present study, we could relate activity of episodic memory circuitry to an AD DPM. Our findings show that neural activity in the episodic memory circuitry changes alongside disease progression and seems to follow MTL atrophy. Consistent with this potential cascade, our longitudinal fMRI findings show that loss of deactivation was most prominent at later disease stages. However, our findings also indicate that the relationship between episodic memory circuit dysfunction and hippocampal volume on the one hand and tau pathology on the other hand is partly independent. This independence indicates that memory function may be associated with multiple processes along the AD cascade including synaptic dysfunction and neurodegeneration. Finally, neural activity in the episodic memory network predicted cognition independent of AD biomarkers and MTL volume.

We started our analysis by first conceptualizing a continuous Bayesian DPM^13^ using longitudinal CSF, volume and cognition data based on the ATN framework. We obtained biomarker progression curves and disease stage scores. The disease stages provide probabilistic information about the position of a participant on an arbitrary disease time frame. We found that our disease time frame was around 20 years (Fig. 1A,B). This is in line with previous conceptualizations showing that AD may unfold over more than two decades^10^. As predicted, disease stage scores were associated with clinical groups, ATN staging and memory accuracy in the fMRI task (A prime) (Fig 1C,D,E). Finally, the time points of fastest change in our data-driven model (Fig. 1B) confirm previously hypothesized trajectory patterns^44^ and results from previous DPM approaches^45^.

After establishing an AD progression marker using the DPM framework, we investigated the relationship between fMRI activations during episodic memory encoding and our DPM scores. Globally, our results indicate a conjoint reduction of encoding-related activations and deactivations as individuals progress along the AD trajectory (Fig. 2B). Thus, individuals in our sample show less pronounced brain activity differences between successfully memorized and later forgotten stimuli as cognition declines, an observation in line with the recently reported risk-dependent reduction of fMRI subsequent memory effects across the Alzheimer’s risk spectrum^39^. Further, our results indicate that the effect sizes for deactivation were larger than for activation, resonating with previous studies that show that task-negative areas overlapping with the DMN^18^ are affected in AD^24,26,30^. Reduced DMN deactivations during episodic encoding can be found even in healthy older adults^35,37^, and they have been associated with relatively poorer memory performance^25,38^.

Given the non-linear course of biomarker abnormalities from our DPM, we next explored potential non-linear associations of regional deactivations and activations (from the fMRI analysis) with disease progression. Thereby, we could identify time points of fastest brain activity change. Our findings suggest that fMRI deactivations may become abnormal before overt cognitive deficits, but after CSF biomarkers and MTL volume (Fig. 2C). Importantly, this was also evident in our longitudinal results, showing more pronounced effects in change of deactivation over time for later-stage participants than for earlier-stage participants (Fig. 4A). This is in contrast to studies suggesting that episodic memory circuitry dysfunction may appear as one of the earliest symptoms in the cascade^46^.

Next, we assessed individual contributions of each DPM marker to activity differences (Fig 3). Our results suggest that synaptic dysfunction and neurodegeneration are independent drivers of episodic memory circuit dysfunction. We found that the emergence of tau was related to a reduction in deactivation and activation, respectively, even when controlling for hippocampal and entorhinal volume. This is in line with a previous study showing that novelty-related activation was related to tau pathology while correcting for hippocampal volume in humans^8^. Moreover, our results resonate with previous studies reporting that tau oligomers can directly lead to synaptic dysfunction (for a review, see^47^), particularly when co-occurring with amyloid pathology^5,8^. These results indicate that deactivation changes in the episodic memory network in AD are partly a consequence of MTL atrophy and partly reflect tau-mediated synaptic dysfunction independent of atrophy. Finally, deactivation changes in episodic memory were related to cognition independently of hippocampal and entorhinal volume, suggesting a relationship between synaptic dysfunction and cognition^7^.

Taken together, our findings provide the first longitudinal evidence about the relationship between episodic memory circuit dysfunction and the Alzheimer’s disease cascade. Our DPM, our analysis of non-linearity, the results from the dependency analysis, and the results from our longitudinal analysis enable us to position episodic memory circuit dysfunction within the ATN framework. As illustrated in Figure 5, our data confirm a neuropathological cascade from amyloid pathology (A), to tau pathology (T), to neurodegeneration (N) and finally to cognitive impairment (C). We suggest that episodic memory circuit dysfunction (E) partly directly follows T and is partly preceded by N. We also suggest that E regularly precedes C. This extended ATN model has implications for the treatment of AD, because it identifies circuit dysfunction as a probable cause for cognitive impairment that is independent of atrophy and therefore potentially reversible. Our extended ATN model thus suggests that there is a time-window within the AD cascade in which a reduction of amyloid and tau pathology can improve cognition, even if there is already some degree of irreversible neurodegeneration.

**Figure 5.**
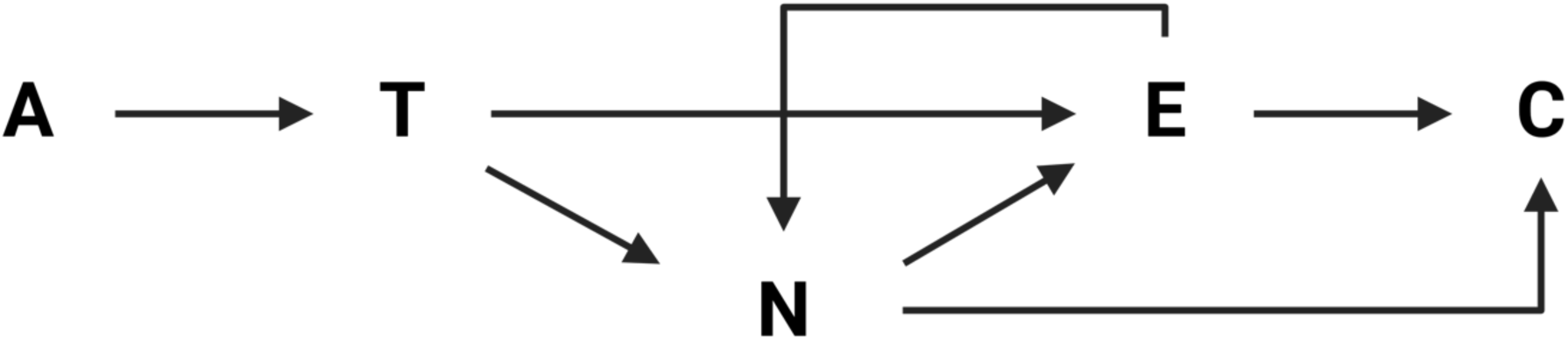
An extended ATN model including episodic memory circuit dysfunction. Our results confirm previous hypothetical conceptualizations of an AD cascade with cognitive impairment (C) being preceded by neurodegeneration (N), tau pathology (T) and amyloid pathology (A). Here, we additionally incorporate episodic memory dysfunction (E) as being partly driven by T and partly by N. E may furthermore result in more N. More explanations in the text.

This study is not without limitations. First, a small number of participants contributed data to the DPM beyond 15 years of estimated disease duration. Due to the low number, and in the light of the previously reported poor expression of fMRI episodic encoding effects in individuals with manifest AD^39,40^, results particularly for the very end of the disease stage spectrum need to be interpreted with caution. Second, we did not correct volumes for age in our DPM approach before entering the DPM algorithm. This was done to preserve the disease-related variability in our sample, since groups of MCI and AD were older than the healthy subsample. Third, we used CSF biomarkers instead of regional biomarker load (PET) resulting in a global estimate for amyloid and tau biomarker status. In particular, a previous study showed that local tau depositions are associated with local changes of activation in episodic memory systems^48^. Fourth, fMRI was not corrected for cerebrovascular contributions in our study^49^. For instance, it was recently shown that variability in cerebrovascular reactivity has an influence on the BOLD response^50^ and thus, on the estimation of effects and, ultimately, the activation values from our first level models. Finally, the fMRI-related activation was characterized by large intra- and interindividual variability, such that our results need to be interpreted with caution. Thus, in order to confirm our extended ATN model, larger samples with longitudinal task-fMRI, CSF measurements, regional biomarker load (PET), and cerebrovascular data are needed. Future research might therefore examine individual change-change associations between activation and local deposition of biomarkers using PET.

In conclusion, this study informed an extended ATN model incorporating episodic memory circuit dysfunction. We found that over the course of the disease, activation and deactivation during successful encoding diminish, partly as a consequence of tau pathology and partly as a consequence of neurodegeneration. These findings help to narrow a knowledge gap of the relationship between AD biomarkers, brain dysfunction and cognition.

## 4. Online Methods

### 4.0 Use of artificial intelligence software

We declare that we used ChatGPT (version 4, https//chat.openai.com/) to aid with grammar and spelling while writing the manuscript. We did not use any other artificial intelligence software for purposes other than the ones declared.

### 4.1 Participants

Participants were part of the multi-center DZNE Longitudinal Cognitive Impairment and Dementia study (DELCODE; Jessen et al., 2018). All participants gave written informed consent in accordance with the Declaration of Helsinki, and the DELCODE study was approved by the ethics committees of all participating institutions. DELCODE was registered with the German Clinical Trials Register (https://www.bfarm.de/EN/BfArM/Tasks/German-Clinical-Trials-Register/_node.html; study ID: DRKS00007966).

In total, 1011 individuals were enrolled in the study. They participated in a range of neuropsychological tests, structural MRI and fMRI (resting-state and/or task-based, diffusion MRI) sessions, as well as cerebrospinal fluid (CSF) biomarker measurements.

Groups were diagnosed based on criteria reported in Jessen et al. (2018). The diagnosis of SCD was given if participants reported subjective memory impairment but scored above the −1.5 standard deviations (SD) cutoff on all subtests of the CERAD-plus test battery adjusted for age, sex, and education. Healthy subjects with above cutoff performance and without any subjective memory impairment were classified as CN. Participants with age-, sex-, and education adjusted memory performance below −1.5 SD on the delayed recall trial of the CERAD-plus episodic memory test were diagnosed with amnestic MCI. Participants were classified as DAT if they fulfilled NINDCS/ADRDA criteria and had a CERAD-plus score of below −1.5 SD, an extended MMSE score between 18-26, and a clinical dementia rating (CDR) greater or equal to one. Additionally, first-degree relatives of patients with a diagnosis of suspected AD (ADrel) were included in DELCODE if their performance was within 1.5 SD in the CERAD-plus test battery.

Among the 1011 participants, we excluded 12 participants who converted to non-amnestic MCI by April 2021, and 23 additional subjects who did not belong to any of the AD-related biomarker groups (see section 4.3). Finally, the group of ADrel was excluded from our analysis, since we did not have any *a priori* hypotheses regarding this subsample (for fMRI activation patterns in this group, see^39,40^). The final analysis sample (n = 493) consisted of 165 cognitively healthy controls (CN), 214 participants with SCD, 82 participants with MCI and 32 participants with DAT, all of whom had available baseline fMRI data. Among these, 218 (44%) participants had at least one available CSF measurement.

### 4.2 Cognition

As cognitive variables, we included the Preclinical Alzheimer’s Cognitive Composite (PACC5)^51^ score and the summary score from the cognitive subscale of the Alzheimer’s disease assessment scale (version 13; ADAS-COG-13)^52^. The PACC5 score is a cognitive composite score reflecting various domains from delayed free and cued recall, results from the Mini-Mental State Examination^53^. The ADAS-COG-13 includes 11 items with tasks related to word recall and recognition, object naming, following commands, constructional and ideational praxis, orientation and comprehension^52^.

### 4.3 CSF-biomarkers

CSF biomarkers were obtained via lumbar puncture performed by trained study physicians. CSF samples were centrifuged, aliquoted, and stored at −80°C for retests, and analyzed with commercially available kits (V-PLEX Aβ Peptide Panel 1 (6E10) Kit (K15200E); Innotest Phospho Tau(181P), 81581, Fujirebio Germany GmbH, Hannover, Germany). CSF biomarker data were available for 218 participants.

For this study, we obtained the Aβ42/40 ratio, pTau181 and total tau (t-tau) values. Those were used (1) to train the disease progression model (see below) and (2) to establish AD-biomarker groups based on the AT(N) criteria ^11^. For this, we used DELCODE-specific cutoff points determined via Gaussian mixture modelling from baseline data in a previous study^54^ (amyloid-negative (A-): Aβ42/40 > 0.08; Amyloid-positive (A+): Aβ42/40 ≤ 0.08; Tau-negative (T-): pTau181 < 73.65; Tau-positive (T+): pTau181 ≥ 73.65). More details on the procedures can be found in^41^.

### 4.4 (f)MRI acquisition

For structural MRI, whole-brain T1-weighted magnetization prepared rapid gradient echo sequence (MPRAGE) images were acquired (TE = 437ms, TR = 2500ms, 7° flip angle, 1mm isotropic) on 3 Tesla Siemens MRI scanner systems. Additionally, coronal T2-weighted turbo spin-echo images of the MTL and hippocampus (TE = 354ms, TR = 3500ms, 120° flip angle, 0.5 × 0.5 × 1.5mm resolution) were obtained. Finally, fMRI was acquired by means of echo planar images (TE = 30ms, TR = 2580ms, 80° flip angle, 3.5mm isotropic).

### 4.5 fMRI task

During the fMRI session, participants were presented with a modified version of an incidental encoding task initially described by Düzel and colleagues ^25^ (for details on the modified version, see^8,26,55^). Participants were presented with 88 novel scenes, evenly split between scenes indoors and outdoors (44 each). Additionally, 44 repetitions of two pre-familiarized scenes – one indoor and one outdoor scene – were presented, each repeated 22 times. The task was administered using Presentation (Neurobehavioral Systems Inc). Participants had to classify each scene as either indoor or outdoor via a button press. Presentation time for each scene was set at 2500 ms. Following a break of 60 min, participants underwent a memory test in which they were instructed to rate their confidence in their recognition of the previously presented new scenes using a 5-point Likert scale.

### 4.6 fMRI preprocessing and first-level modelling

Functional MRI preprocessing was done in SPM12 (Welcome Trust Centre for Human Neuroimaging, UCL, UK). The functional MRIs were slice-time corrected, unwarped, realigned, segmented, coregistered with the structural images, normalized to a population standard space via geodesic shooting, normalized to MNI space via an affine transformation and, finally, spatially smoothed with a Gaussian filter kernel at 6 mm full width half maximum (FWHM).

For the first-level general linear models (GLMs), the canonical hemodynamic response function with a 128-second high-pass filter and no global scaling was used (see^26^). Regressors of interest were the novelty regressor (all novel images), a successful memory regressor obtained from a parametric arcsine transformation of the novelty regressor modulated with the confidence ratings from the recognition task^39^, and a regressor for the familiar images. Additionally, each GLM included the six rigid-body motion parameters determined from realignment as covariates of no interest, plus a single constant representing the implicit baseline. The contrast of interest was the successful memory encoding contrast (i.e., the parametric modulation of the novelty regressor with recognition confidence in the subsequent memory test).

### 4.7 Disease progression model for staging based on ATN pathology

For quantification of each patient’s disease stage during progression towards AD, we utilized a continuous multivariate DPM based on Bayesian Gaussian Process (GP) regression ^13,56^ (code available from https://gitlab.inria.fr/epione/GP_progression_model_V2). In this approach introduced by Lorenzi et al.^13^, GPs were used to empirically describe smooth monotonically increasing transition curves from normal to abnormal biomarker levels as proposed in REF^44^ (see REF^13^ for mathematical details). Our DPM application in this study focused on the following groups of variables based on the ATN pathological classification system and cognitive performance: (A) Biomarkers from CSF were Aβ42/40 ratio and pTau181 levels. (B) Atrophy measures comprised bilateral hippocampus and entorhinal cortex volumes from FreeSurfer (version 7, longitudinal pipeline, corrected for total intracranial volume (TICV)). (C) Cognition was operationalized in terms of the PACC5 and the ADAS-COG-13 summary scores. Importantly, the model inversion based on observation of those features (including available follow-ups) estimates the most probable individual patient’s disease stage continuously (including its uncertainty) and a data-driven disease progression curve for each of the fitted biomarkers.

#### 4.7.1 Participant selection for the model fit

For DPM fit, we only included DELCODE participants who had at least one complete measurement occasion of biomarkers^13^. Out of those, we excluded participants with a biomarker profile outside the Alzheimer’s continuum (e.g., A-T+; see^57^). This was further refined by restricting the clinical AD-risk groups (i.e., SCD, MCI, and DAT) to those with amyloid positivity (A+). Additionally, we used clinical conversion data to exclude participants who later converted to non-amnestic MCI or to non-Alzheimer’s type dementia such as dementia with Lewy bodies, Parkinson’s disease dementia, semantic dementia, or vascular dementia. This resulted in a subsample of 208 participants (80 CN, 57 A+ SCD, 44 amnestic A+ MCI, 27 A+ DAT). Further details on the descriptive statistics of the model estimation sample and parameter settings, as well as schematic representation can be found in the Supplementary Table 2 and Supplementary Figure 6, respectively. After model fitting, model inversion was done on the remaining participants who were not part of the fitting process.

### 4.8 Statistical analyses

#### 4.8.1 Demographics and exploratory analyses

All statistical analyses (except for image-based analyses) were carried out in R (version 4.2.1). Significance levels were set at *p* < 0.05. First, group differences in demographics were analyzed by one-way analyses of variance (ANOVA). Wherever normality was violated, non-parametric Kruskal-Wallis tests were used. Group differences in sex were analyzed by means of chi-square tests. *Post-hoc* group differences were analyzed by two-sample t-tests or Wilcoxon rank-sum tests and corrected for multiple testing by using the Bonferroni-Holm correction. Relationships between disease stage and fMRI task performance were analyzed with bivariate partial correlations adjusting the variables for age, sex, education, and scanning site. Individual associations between biomarkers and disease stage were analyzed using Pearson and Spearman correlations. Next, to assess the association between disease stages and diagnostic groups and AT biomarker categories, respectively, we calculated separate one-way analyses of covariance (ANCOVAs) with disease stage as the dependent variable and diagnostic group and AT biomarker status as independent variables, adjusting for age, sex, education and MRI scanning site.

#### 4.8.2 Voxel-wise image analyses using the Sandwich estimator toolbox (SwE)

After establishing a marker of disease progression for AD, we were interested in the association of that marker with the fMRI activation contrasts on the voxel-level. We expected a region-specific association of activation with disease progression. The analysis was carried out using the Sandwich estimator toolbox^58^ (v2.2.2, MATLAB). All available longitudinal scans were included in the fitting process to obtain between-subjects effects and additionally estimate longitudinal effects of time. The design matrix included an intercept, linear and quadratic terms of the disease stage, intraindividual time coded as difference from the intraindividual mean timepoint in years, age, sex, education, and the interaction terms time*covariates. For inference, we used the Wild Bootstrap method^59^ with 1000 repetitions. T-contrasts for the effects of interest (linear and quadratic effects of disease stage) were obtained and thresholded using family-wise error (FWE) correction with *p* < 0.05.

#### 4.8.3 Estimation of fMRI-based activity curves during disease progression

Next, we next were interested in the progression patterns of activation and deactivation over the disease stages. To do so, we extracted mean activation and deactivation values from the FWE-thresholded (de-)activation maps for each of the available measurement time points. These were submitted to smoothing splines using the ss() function from the *npreg* package in R ^60^. The independent variable was the disease stage obtained from DPM. The dependent variable was the average activation or deactivation value. In addition to the function’s default values, number of knots were *a-priori* set to 3. To estimate the time point of fastest activation change over the disease stages, we additionally calculated first derivatives of the cluster-level smoothing splines over disease stage using the *pracma* package in R^61^. This provided us with an estimate of change of episodic memory-related deactivation and activation at each point along the disease trajectory line. This was done to compare the relative change magnitudes of markers from the DPM to the change magnitudes of episodic memory activation.

#### 4.8.4 Individual contributions of DPM markers to episodic memory activity differences

Next, we were interested in the respective contributions of DPM markers to activity differences. For this, we first calculated two separate one-way ANCOVAs for the association between episodic memory deactivation and activation and AT staging groups while controlling for hippocampal and entorhinal volume and demographics age, sex, and education. Second, we were interested in the association between volumes and activity while controlling for CSF biomarkers. For this, we calculated semi-partial Pearson correlations between episodic memory activity and volumes adjusted for demographics and CSF-Aβ42/40 ratio and CSF-pTau181. Finally, to check the association between activity and cognition, we calculated semi-partial Pearson correlations adjusting cognition for demographics and hippocampal and entorhinal volume.

#### 4.8.5 Longitudinal activation changes over time

Lastly, we aimed at testing above findings longitudinally. First, we calculated linear mixed effect models (LMEs) in R with episodic memory activity as the dependent variable. The effect of interest was the interaction between time and disease stage. Here, disease stage was split into two groups based on the median disease stage in the sample. Covariates were the interactions between time and demographics age, sex and education. We calculated LMEs with and without participant ID as random slope effect, but the former ones did not converge, so the latter ones were used for the analysis. Second, we were interested if longitudinal activation change over follow-ups would align with the actual intra-individual stage progress of an individual over follow-ups (as indicated by ATN and Cog). For this, we obtained disease stage estimates for each measurement occasion independently for each participant using the available data. Next, we used linear regression to estimate rates of change along the disease progression with longitudinal disease stage estimates as the dependent variable and intra-individual time as the independent variable. Additionally, we repeated this for longitudinal mean activation values by submitting the activity values as the dependent variable and intra-individual time as the independent variable to simple linear regressions. Finally, we extracted the beta coefficients from both simple linear regressions and correlated them using Pearson correlation. We expected that higher change in contrast value would be related to a faster disease progression.

## Supplementary material to

**Supplementary Table 1.** Baseline characteristics of the final analysis sample (n = 493)

|  | Missing | CN (n = 165) | SCD (n = 214) | MCI (n = 82) | DAT (n = 32) | Group test P-value | Pairwise comparisons |
| --- | --- | --- | --- | --- | --- | --- | --- |
| Age (y) | - | 69.17 (5.24) | 70.51 (6.05) | 72.72 (5.65) | 73.86 (6.26) | < 0.001 | CN vs. SCD, MCI, DAT<br>SCD vs. MCI, DAT |
| Sex (% female) | - | 105 (63.64) | 104 (48.60) | 43 (52.44) | 18 (56.25) | 0.041 | CN vs. SCD |
| Education (y) | - | 14.61 (2.68) | 15.28 (2.93) | 13.90 (3.02) | 13.09 (2.86) | < 0.001 | CN vs. DAT<br>SCD vs. MCI, DAT |
| APOE ε4 carrier (%) | 5 (1.01) | 38 (23.60) | 70 (32.86) | 41 (50.00) | 21 (65.63) | < 0.001 | CN vs. MCI, DAT<br>SCD vs. MCI, DAT |
| CSF-Aβ42/40 ratio | 275 (55.78) | 0.10 (0.02) | 0.09 (0.03) | 0.07 (0.03) | 0.05 (0.02) | < 0.001<sup>†</sup> | CN vs. MCI, DAT<br>SCD vs. MCI, DAT<br>MCI vs. DAT |
| CSF-pTau<sub>181</sub> (pg/ml) | 275 (55.78) | 45.86 (13.51) | 51.68 (20.50) | 68.02 (34.00) | 99.06 (42.55) | < 0.001<sup>†</sup> | CN vs. MCI, DAT<br>SCD vs. MCI, DAT<br>MCI vs. DAT |

The groups were significantly different in terms of age ( $F_{3,489} = 10.59, p < .001$ ), sex ( $\chi^2_3 = 8.29, p = .041$ ), years of education ( $F_{3,489} = 8.45; p < .001$ ) and APOE ε4 carriership ( $\chi^2_3 = 31.26, p < .001$ ). Additionally, CSF-Aβ42/40 ratios ( $F_{3,76,297} = 52.17, p < .001$ ) and CSF-pTau<sub>181</sub> levels ( $F_{3,62,296} = 14.60, p < .001$ ) were different across the groups. CN = healthy controls, SCD = subjective cognitive decline, MCI = mild cognitive impairment, DAT = mild dementia of the Alzheimer's disease type. <sup>†</sup>F-test calculated using the Welch method. Pairwise comparisons denote significant pairwise post-hoc comparisons after correction using Bonferroni-Holm.

**Supplementary Table 2.**
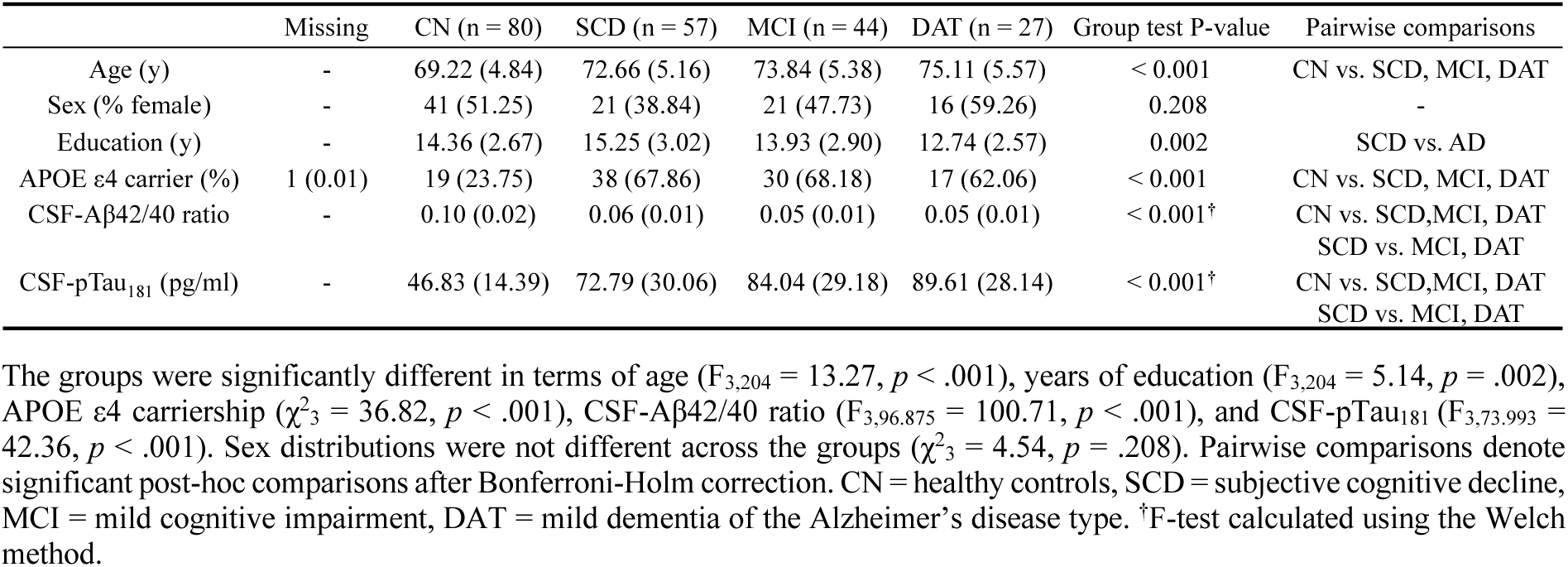
Baseline characteristics of the model training sample (n = 208)

**Supplementary Table 3.** *Post-hoc* comparisons for the association between disease stage and diagnostic groups.

| Group Comparison | Mean Difference | SE | P<sub>uncorr</sub> | P<sub>corr</sub> |
| --- | --- | --- | --- | --- |
| SCD > CN | 1.53 | 0.359 | < 0.001 | < 0.001 |
| MCI > CN | 5.85 | 0.694 | < 0.001 | < 0.001 |
| DAT > CN | 12.91 | 0.755 | < 0.001 | < 0.001 |
| MCI > SCD | 4.32 | 0.686 | < 0.001 | < 0.001 |
| DAT > SCD | 11.38 | 0.747 | < 0.001 | < 0.001 |
| DAT > MCI | 7.06 | 0.954 | < 0.001 | < 0.001 |

Pairwise comparisons from the ANOVA assessing the association between diagnostic groups and disease stage. P<sub>uncorr</sub> = raw p-values. P<sub>corr</sub> = corrected p-values using Bonferroni-Holm. CN = healthy controls, SCD = subjective cognitive decline, MCI = mild cognitive impairment, DAT = mild dementia of the Alzheimer's disease type.

**Supplementary Table 4.** *Post-hoc* comparisons for the association between disease stage and AT staging groups.

| Group Comparison | Mean Difference | SE | P<sub>uncorr</sub> | P<sub>corr</sub> |
| --- | --- | --- | --- | --- |
| A+T- > A-T- | 3.27 | 0.733 | < 0.001 | < 0.001 |
| A+T+ > A-T- | 9.1 | 0.716 | < 0.001 | < 0.001 |
| A+T+ > A+T- | 5.83 | 0.916 | < 0.001 | < 0.001 |

Pairwise comparisons from the ANOVA assessing the association between AT staging groups and disease stage. P<sub>uncorr</sub> = raw p-values. P<sub>corr</sub> = corrected p-values using Bonferroni-Holm. A-T- = amyloid and tau negative, A+T- = amyloid positive and tau negative, A+T+ = amyloid and tau positive.

**Supplementary Table 5.**
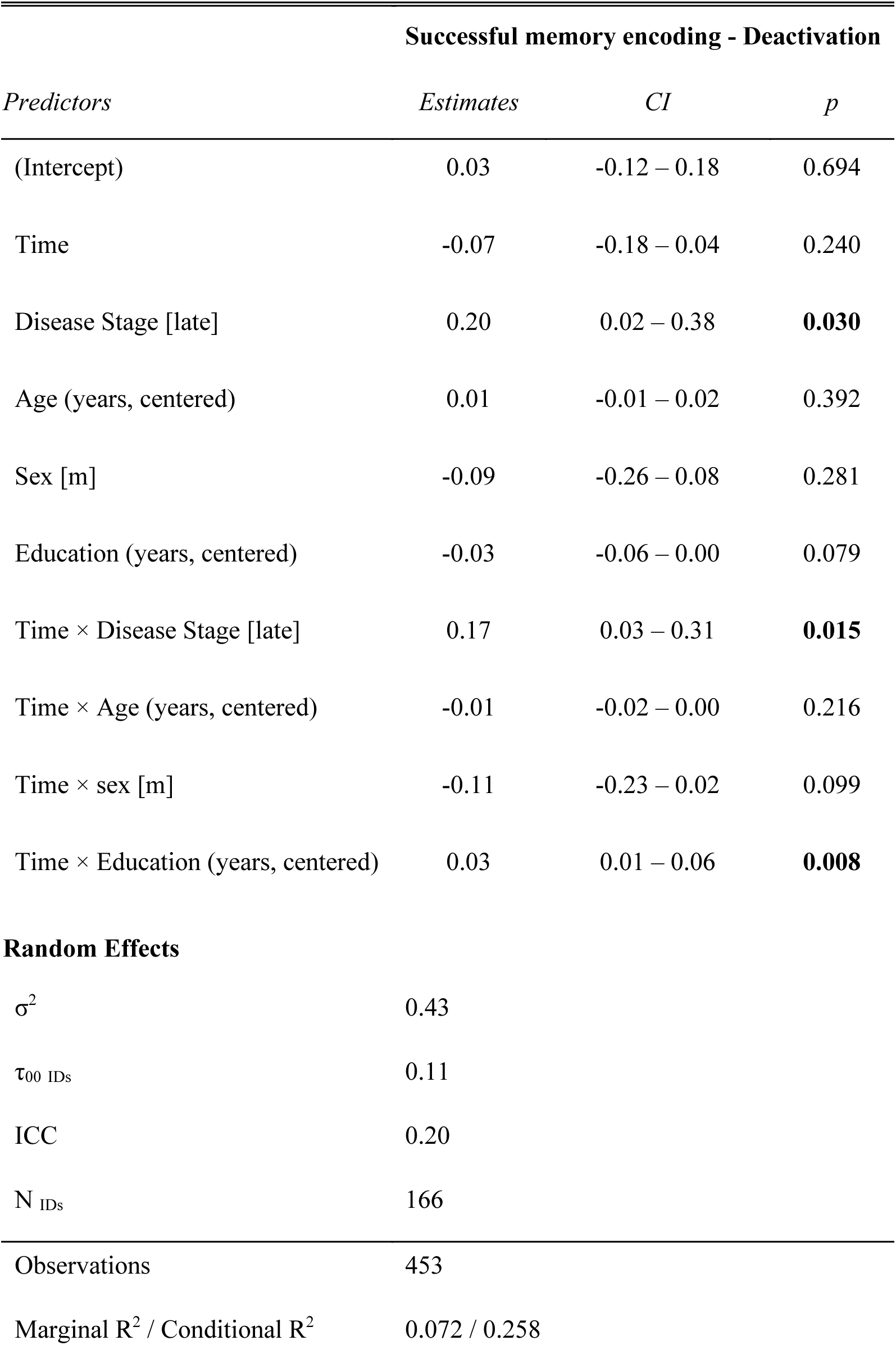
Linear mixed effects model output for the association between successful memory encoding-related deactivation and disease stage.

| Successful memory encoding - Deactivation |  |  |  |
| --- | --- | --- | --- |
| *Predictors* | *Estimates* | *CI* | *p* |
| (Intercept) | 0.03 | -0.12 – 0.18 | 0.694 |
| Time | -0.07 | -0.18 – 0.04 | 0.240 |
| Disease Stage [late] | 0.20 | 0.02 – 0.38 | **0.030** |
| Age (years, centered) | 0.01 | -0.01 – 0.02 | 0.392 |
| Sex [m] | -0.09 | -0.26 – 0.08 | 0.281 |
| Education (years, centered) | -0.03 | -0.06 – 0.00 | 0.079 |
| Time × Disease Stage [late] | 0.17 | 0.03 – 0.31 | **0.015** |
| Time × Age (years, centered) | -0.01 | -0.02 – 0.00 | 0.216 |
| Time × sex [m] | -0.11 | -0.23 – 0.02 | 0.099 |
| Time × Education (years, centered) | 0.03 | 0.01 – 0.06 | **0.008** |
| **Random Effects** |  |  |  |
| $\sigma^2$ | 0.43 | | |
| $\tau_{00}$ IDs | 0.11 | | |
| ICC | 0.20 |  |  |
| N IDs | 166 |  |  |
| Observations | 453 |  |  |
| Marginal R<sup>2</sup> / Conditional R<sup>2</sup> | 0.072 / 0.258 |  |  |

**Supplementary Table 6.** Linear mixed effects model output for the association between successful memory encoding-related activation and disease stage.

| Successful memory encoding - Activation |  |  |  |
| --- | --- | --- | --- |
| *Predictors* | *Estimates* | *CI* | *p* |
| (Intercept) | 0.18 | 0.01 – 0.35 | **0.037** |
| Time | -0.05 | -0.16 – 0.06 | 0.400 |
| Disease Stage [late] | -0.24 | -0.45 – -0.04 | **0.018** |
| Age (years, centered) | -0.03 | -0.05 – -0.01 | **0.004** |
| Sex [m] | -0.05 | -0.24 – 0.14 | 0.603 |
| Education (years, centered) | 0.06 | 0.02 – 0.09 | **0.001** |
| Time × Disease Stage [late] | 0.12 | -0.02 – 0.26 | 0.098 |
| Time × Age (years, centered) | 0.00 | -0.01 – 0.01 | 0.922 |
| Time × Sex [m] | -0.03 | -0.16 – 0.10 | 0.656 |
| Time × Education (years, centered) | 0.01 | -0.01 – 0.03 | 0.409 |
| **Random Effects** |  |  |  |
| $\sigma^2$ | 0.46 | | |
| $\tau_{00}$ IDs | 0.16 | | |
| ICC | 0.26 |  |  |
| N<sub>IDs</sub> | 166 |  |  |
| Observations | 459 |  |  |
| Marginal R<sup>2</sup> / Conditional R<sup>2</sup> | 0.129 / 0.357 |  |  |

**Supplementary Figure 1.**
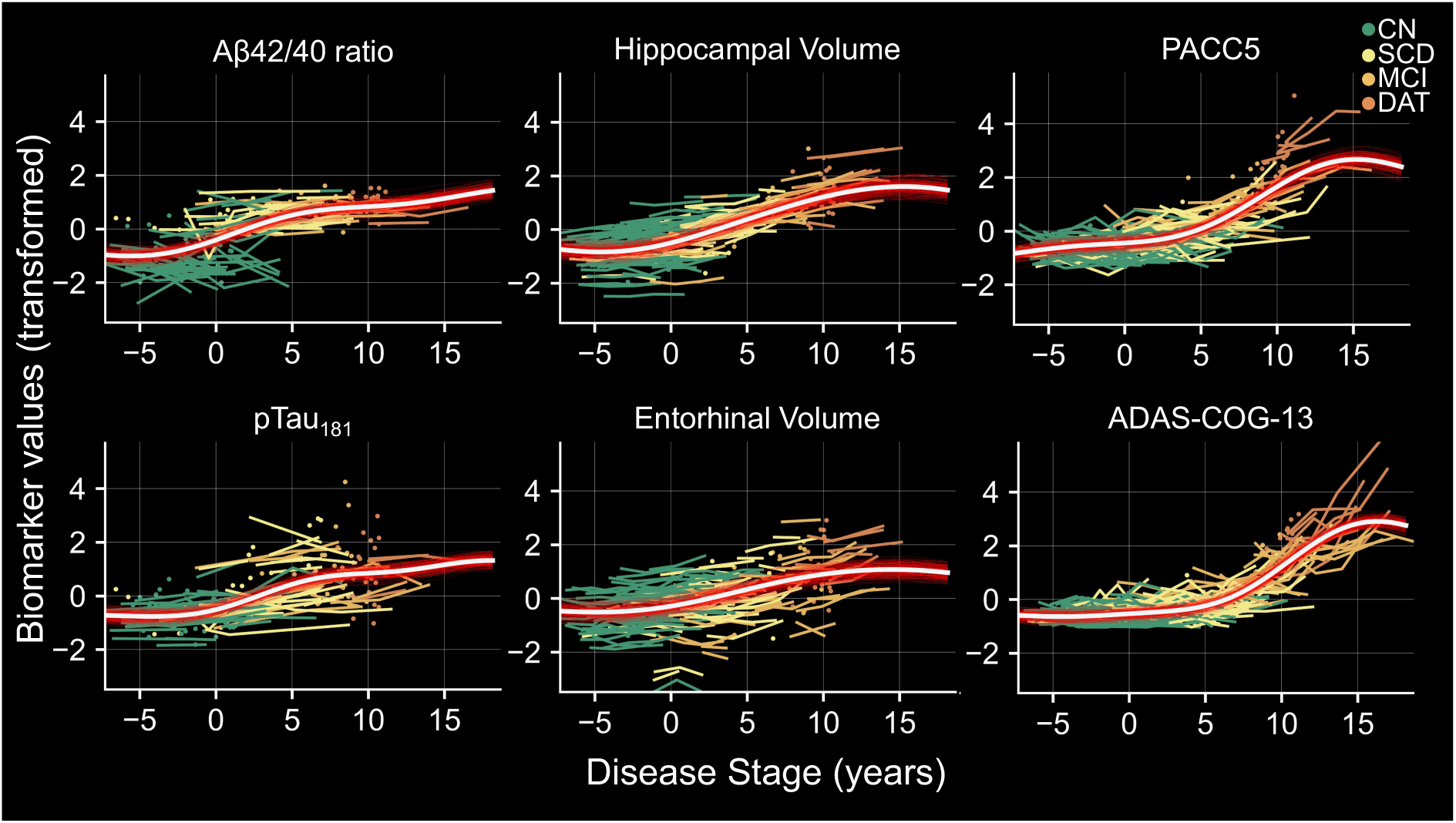
Model-based Gaussian-Process realizations from the model posterior distribution. Red curves show 200 individually sampled realizations from the model posterior. Curves in white show the average GP. Colored lines indicate 739 longitudinal data points from the 208 participants added into the model. All available longitudinal data was used. CN = healthy controls, SCD = subjective cognitive decline, MCI = mild cognitive impairment, DAT = mild dementia of the Alzheimer‘s disease type.

**Supplementary Figure 2.**
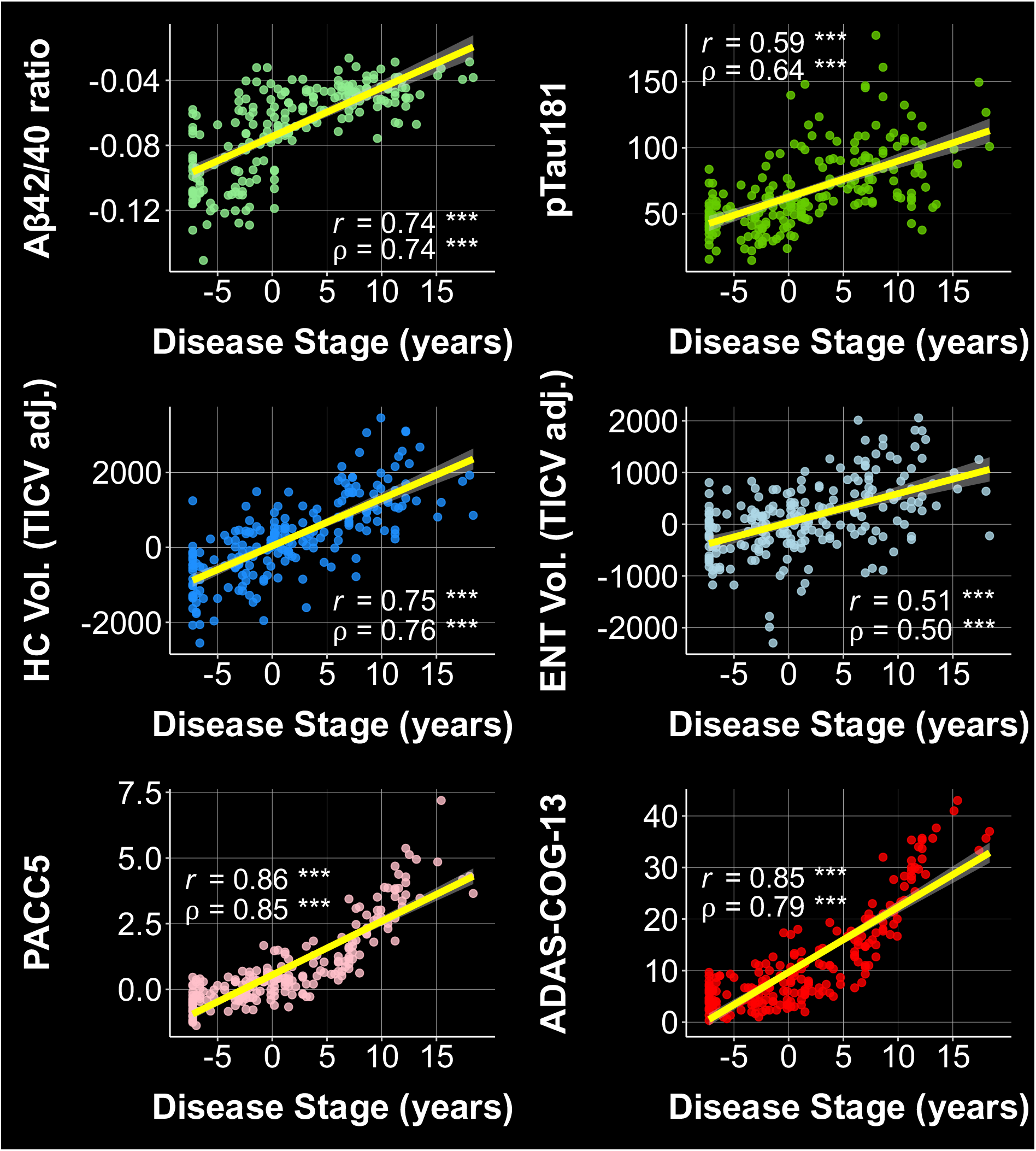
Bivariate correlations between variables used in the DPM and the DPM-derived disease stage for the model fitting sample (n = 208). Here, we report Pearson (r) and Spearman rank (ρ) correlations for each variable in the fitting sample comprising n_CN_ = 80, n_SCD_ = 57, n_MCI_ = 44, n_DAT_ = 27 participants. Note that the CSF ratio, MTL volumes and PACC5 were multiplied with −1. Therefore, an increase in disease stage is related to an increase in more pathological DPM marker values. HC Vol.: Hippocampal Volume, ENT Vol.: Entorhinal Cortex Volume. Both volumes were adjusted for total intracranial volume (TICV). *p* < .001***, *p* < .01**, *p* < .05*, *p* > .05^n.s.^

**Supplementary Figure 6.**
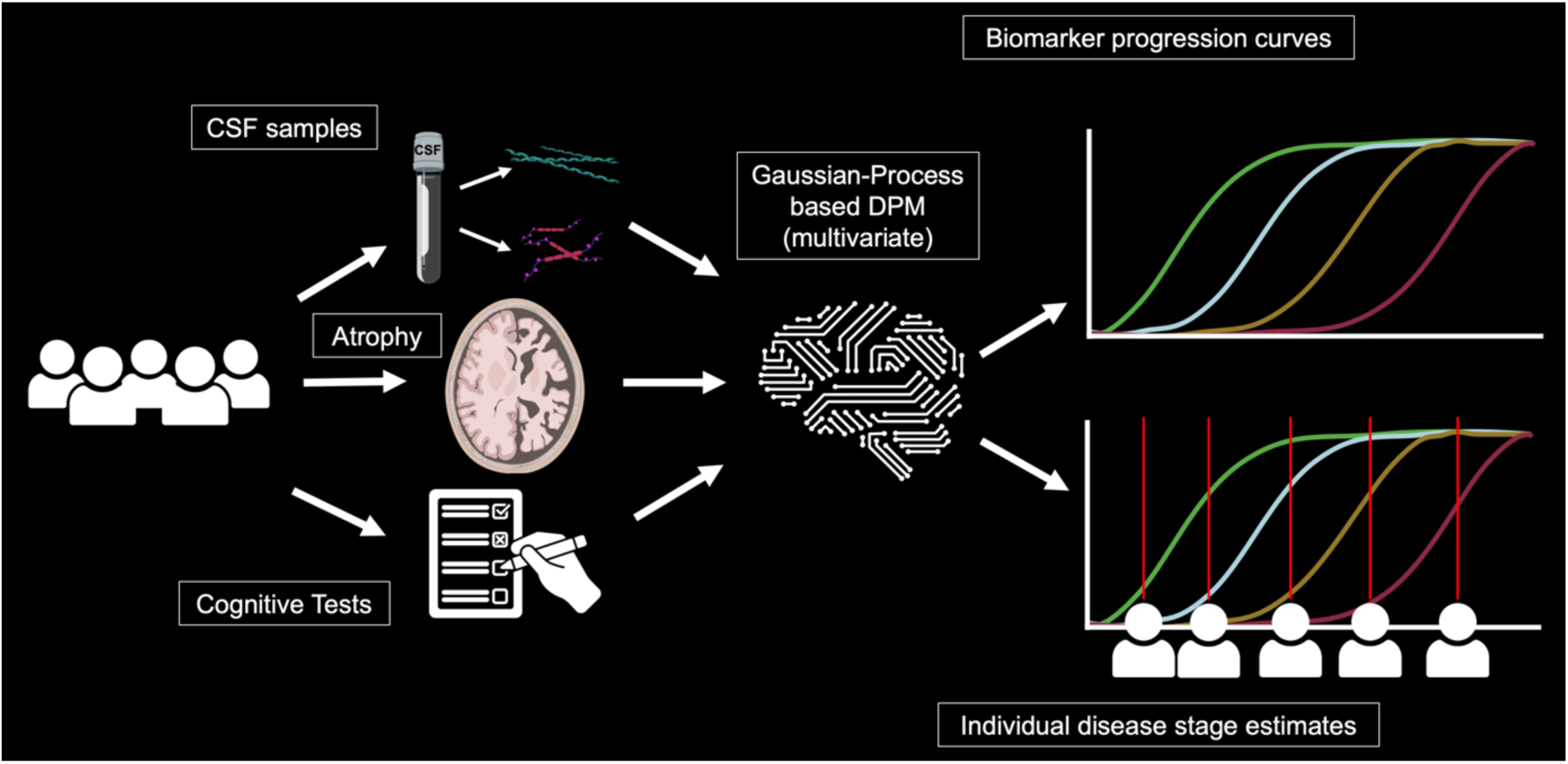
Schematic representation of analyses in our study. For model fit, we selected 208 participants with at least one complete measurement occasion of biomarkers (see Lorenzi et al., 2019). If available, we also included longitudinal biomarker information, such that 739 data points were included in the model training procedure. Model training parameters were kept at default values: Outer iterations = 6; Inner iterations = 200; trade-off = 100. The trade-off describes the strength of the monotonicity constraint in relation to the model fit. Monotonicity constraint for each biomarker was set to 1. This was done such that increasing levels in biomarkers would be associated with higher burdens in all respective domains. For this, the CSF-Aβ42/40 ratio and volume values were multiplied by −1. After model fitting, the disease stage estimates for all participants were obtained using the *Predict* method. Lastly, time was operationalized as time-since-baseline expressed in years. All variables were z-standardized prior to model fitting. Figure created with BioRender.com

## Notes

### Competing Interest Statement

The authors have declared no competing interest.

